# Percentage-Based Author Contribution Index. A universal, measure of author contribution to scientific articles

**DOI:** 10.1101/138875

**Authors:** Stéphane Boyer, Takayoshi Ikeda, Marie-Caroline Lefort, Jagoba Malumbres-Olarte, Jason M. Schmidt

## Abstract

Deciphering the amount of work provided by different co-authors of a scientific paper has been a recurrent problem in science. Despite the myriad of metrics available, the scientific community still largely relies on the position in the list of authors to evaluate contributions, a metric that attributes subjective and unfounded credit to co-authors. We propose an easy to apply, fair and universally comparable metric to measure and report co-authors contribution in the scientific literature. The proposed Author Contribution Index (ACI) is based on contribution percentages provided by the authors, preferably at the time of submission. Researchers can use ACI for a number of purposes, including comparing the contributions of different authors, describing the contribution profile of a researcher or analysing how contribution changes through time. We provide an example analysis based on data collected from 97 scientists from the field of ecology who voluntarily responded to an online anonymous survey.

## Introduction

Deciphering the role and quantifying the amount of work provided by different co-authors of a particular paper has been a recurrent problem for the scientific community [2, 3, 16]. The position in the list of authors is commonly used to infer co-authors’ contribution and a number of systems have been proposed on this basis. They range from simple calculations based on the rank of the authors such as harmonic authorship credit, fractional authorship credit, inflated authorship [1] to more complex credits (e.g. [7]), some even taking into account the controversial journal’s impact factor [16]. However, these metrics are essentially ‘one fits all’ approaches that assume the contribution of each author based on their position in the author list and attributes subjective and unfounded values to these positions. As such they do not attempt to represent and quantify ‘true’ contribution. Despite the growing interest in resolving the issue of authorship contributions in scientific disciplines [1, 3, 15], no standard rank system has been widely recognised or adopted by scientific journals. With this lack of consensus, some journals have implemented a compulsory or recommended section about authors’ contribution. A review of the top 150 ecology journals referenced in ISI Web Of Knowledge revealed that 13.3% of them have adopted this practice (Supporting Information 1). Authors are usually asked to briefly describe which task was conducted by which co-author. Although this information is valuable, it does not provide an objective, straightforward and universal measure of author contribution. For example ‘data collection’ for a review article may simply involve searching a database using specific key words, while it may be a very time consuming task in field ecology, and a highly technical task in computational ecology. So ‘data collection’ can mean very different things depending on the field of study or the type of paper. In addition, individual tasks are often conducted by multiple authors but there is no way of knowing whether one author has contributed more to them. Although some systems propose graded contributions for each task (e.g. lead, equal, supporting role in the CRedIT system), the lack of a continuous value means these systems lack accuracy and it is very difficult to analyse or compare contributions across multiple articles or years. The second common limitation is the complexity of the proposed systems which often deters authors from providing the data and hinders the understanding and use of these data by others. A third major issue is the lack of fairness where often the lead or corresponding author can unilaterally decide on the order of the co-authors and the description of their contribution.

To address these shortcomings, we propose an easy to apply, universally comparable and fair tool to measure and report author contribution.

## A simple and accurate measure: percentage contributions

Percentages are straightforward and can be universally applied independent of research field, the number of co-authors or the nature of the paper (e.g. experimental, review, perspective etc.). Because the authors of a paper are the best placed to make a judgment call about the value of each contribution, it is essential that percentage contributions are determined by authors rather than by a model based solely on the authors’ rank. Although disagreement may occur between co-authors, clarifying contribution among co-authors in the early stages of the research is likely to ease potential tension [12], and in some cases prompt ‘real collaboration’. A possible starting point is to divide 100% by the number of authors and then estimate whether and to what extent each author provided more or less work than the others.

The use of author-provided percentages has been proposed before to reflect the contribution of co-authors accurately (e.g. [16]), but with limited guidance about how to implement it. Verhagen et al. [17] proposed the Quantitative Uniform Authorship Declaration (QUAD) approach, where each author is attributed percentage contributions in four categories: Conception and design, data collection, data analysis and conclusion, manuscript preparation. More recently, a very similar approach was proposed based on scores rather than percentages with the more specific aim of deciding which contributor deserves authorship and which does not [18]. Clement [11] also suggests the use of four categories, albeit slightly different ones (ideas, work, writing, and stewardship). However, an overly complicated metric is likely to deter authors from applying it, and the proposed criteria and categories may not be consistent or have comparable importance across research fields and may not be applicable to every type of article. In addition, many authors suggest that contributions should be restricted to an arbitrary threshold, for example 50% of the average contribution [11], 10% of the total work [17] or a threshold chosen by the authors [18]. Such limitation is likely to introduce major inconsistencies between papers, journals and fields of research, thereby preventing comparison. In addition, these thresholds limit the number of co-authors, which may affect interdisciplinary research and act as incentives to leave out minor contributors, potentially increasing ghost authorship (i.e. the omission of collaborators who did contribute to the work).

We propose that the contribution of each co-author be summarised in one number which must be more than 0% and less than 100% in multiple-authored papers. This provides a metric that is simpler for authors to determine and for the readers to grasp. In addition, this single metric imposes no upper limit on the number of authors. The percentage contribution should be displayed on the published paper either as raw numbers or as a figure (Fig. 1).

**Figure 1.**
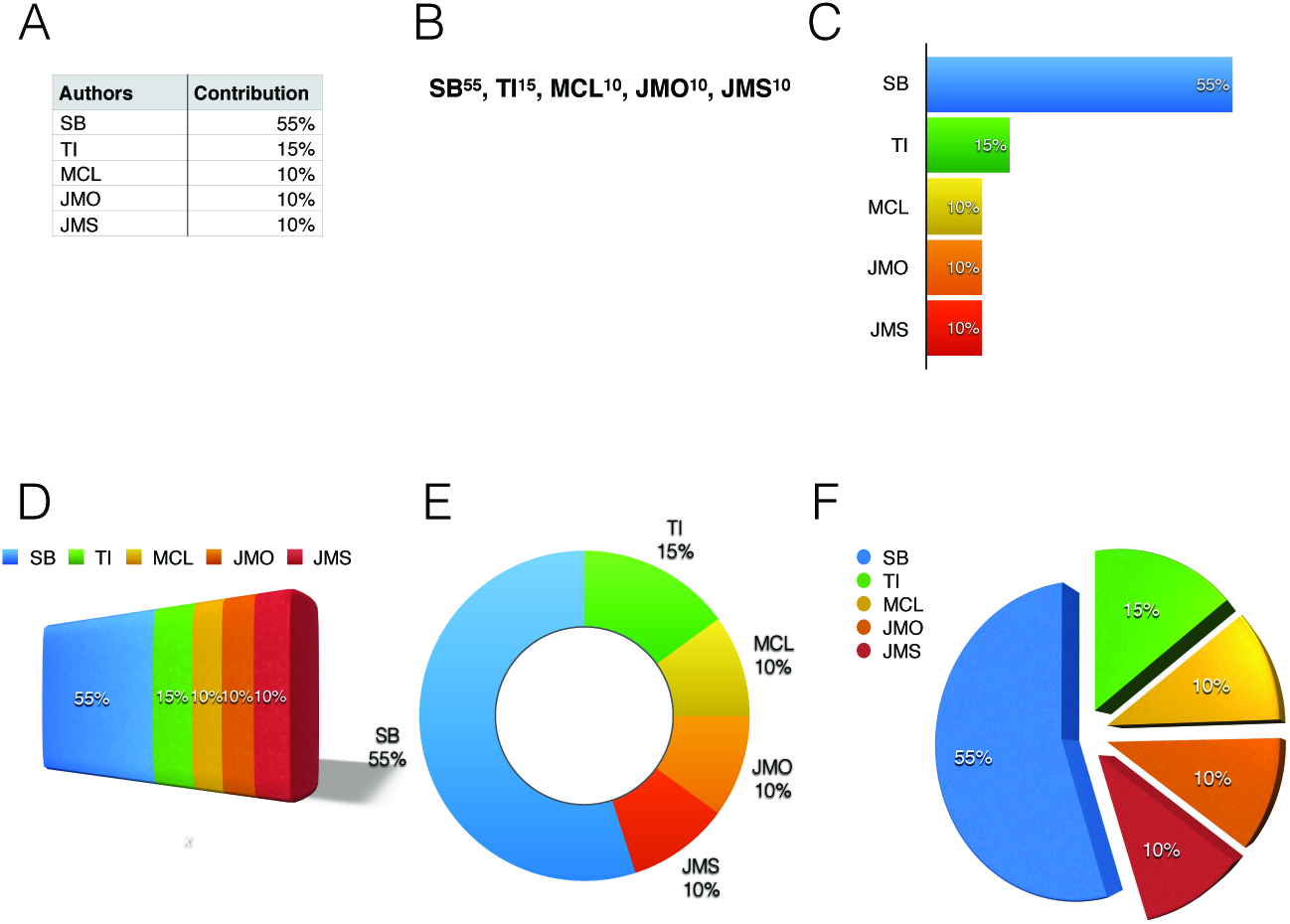
Examples of a table (A), text (B) or figures (C, D, E, F) that could be displayed on published articles to illustrate author contribution percentages. Data correspond to author contributions for the current paper.

We propose that co-authors discuss and agree on their respective contributions prior to submitting their manuscript and these figures be provided by the corresponding author at the submission stage. By confirming their authorship, all co-authors confirm their agreement with their contributions and that of all other authors. This ensures that every published paper displays percentage contributions that have been discussed and agreed upon by every co-author.

## A universally comparable metric: percentage-based author contribution index (ACI)

An outstanding limitation of percentage contributions is that they are difficult to compare across different papers because with more co-authors, it is mathematically more difficult to obtain high percentages. As a consequence, author contributions cannot be directly compared between articles with various number of authors. To allow such comparison, we propose a universal metric that takes into account the number of co-authors: the Author Contribution Index (ACI), calculated from the percentage contribution as per Equation (1).

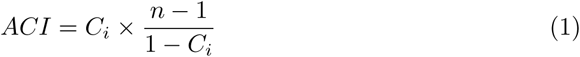

Where for author *i*:

*C_i_* = contribution of author *i* in percentage (must be >0 and <1)
*n* = total number of authors including *i* (must be >1)

ACI reflects the contribution of author as compared to the average contribution of all other authors. It is superior to one when the author’s contribution is larger than the average contribution of all other authors, and inferior to one when the author’s contribution is less than the average contribution of all other authors. For example, on a paper written by three authors, where author *i* contributed 60% of the paper, *ACI_i_* = 3, meaning that author *i* contributed three times more than what the other authors contributed on average. Another useful metric is *log*_10_ (*ACI*), which is positive when the author’s contribution is larger than the average contribution of all other authors, and negative when the author’s contribution is less than the average. This metric is useful to normalise data for further comparison and statistical analyses.

The graph in Fig. 2 displays the universe of all possible ACIs for papers written by up to 200 co-authors. The contribution profile of a particular author can be displayed in the universe of possible ACIs by adding dots, each representing one paper from the author being analysed. From these data, author profiles may appear according to a variety of criteria such as time, author’s seniority, area of research and type of institution where the author works, among others. Based on Equation (1), it is also possible to calculate average ACI for an individual author or to plot the ACI frequency distribution of an individual author based on all or specific parts of his publications.

ACI increases with the proportion of work produced but also with the number of ‘minor’ co-authors (Fig. 2). By giving more weight to main contributors of papers with many co-authors, ACI recognises the skills required and work involved in leading large collaborative projects. Fig. 3 provides examples of how ACIs could be displayed in a paper.

**Figure 2.**
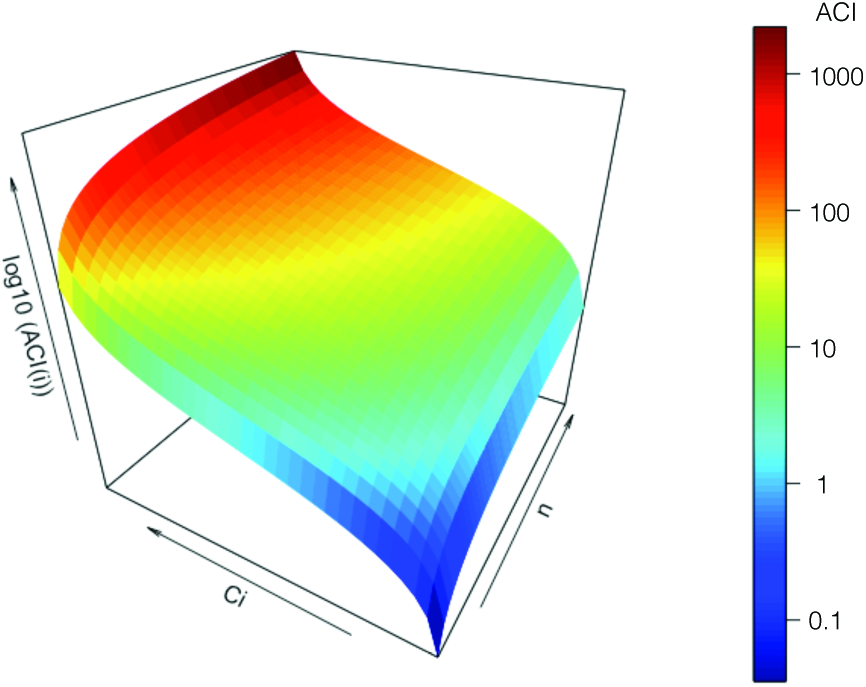
Universe of possible ACIs. X-axis: total number of authors including (n = 2 to n = 200); y-axis: percentage contribution of author (=0.001 to = 0.999); z-axis: author contribution index for author (see Equation (1)). Colours correspond to the value of ACI (see coloured scale on the right).

**Figure 3.**
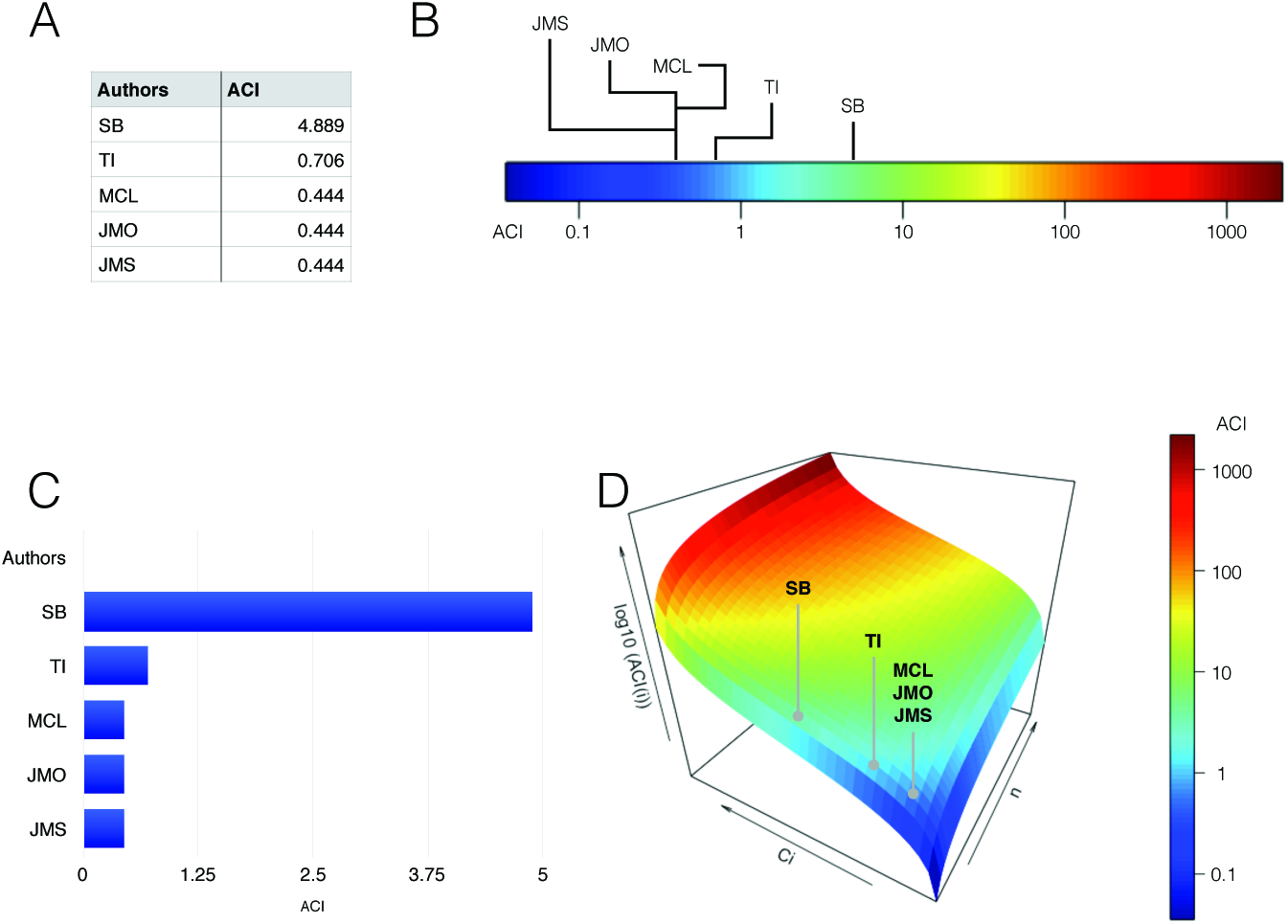
Examples of table (A) or figures (B, C, D) that could be used to display the author contribution index (ACI) for a given paper. Data corresponds to author contributions for the current paper.

## A fair tool: assisting job seekers, recruiters and performance-based evaluations

The scientific community seems to have reached the consensus that journal impact factors are not an accurate measure of the value of a particular article or the value of its author(s) [8]. One of the main reasons is that a very highly-ranked journal may publish few articles that are heavily cited, but it may also publish a large number of papers that will have very little impact. In recent years, article-based impact has been preferred to journal impact factor. For example, the Hirsch index (*h*-index), which is based on the number of citations of one’s papers is now widely used to gauge the output of a scientist. However, the *h*-index can also be manipulated [13] and it does not provide a measure of the amount of work produced by each co-author, which means guest authorship (i.e. inclusion of authors who did not contribute to the work), cannot be accounted for.

Getting a clear idea of the amount of work a scientist is actually providing is difficult if one needs to read through all the authors contribution sections and weigh in the topic, the type of paper, the number of co-authors etc. ACI can provide valuable information for performance-based evaluation processes and could be implemented in existing reporting systems. This includes internal evaluation for career advancement, as well as research productivity evaluation for funding purposes and national-scale ranking schemes (such as the Performance-based Research Fund (PBRF) system in New Zealand or the Research Excellence Framework (REF) in the UK). It is also in the advantage of the candidate to be able to demonstrate his/her actual contribution to a potential recruiter who may ask ‘what have you done on all these papers listed on your CV?’. One could answer such a question by analysing the distribution of a scientist’s ACI and its evolution through time or by calculating and comparing his/her average ACIs in experimental, review and perspective papers. ACI could also be used as an additional metric in network-based collaboration analyses (eg. [10]) or to further inform composite citation indicators (e.g. [6]).

## A first look: testing ACI

Here we provide an analysis of ACI calculated across the past 3 years (January 2014-December 2016) for 97 ecologists from 19 different countries. ACIs were calculated based on contribution percentages provided by scientists who volunteered to respond to an online survey (Supporting Information 2). Respondents comprised postgraduate students, postdoctoral fellows, early-career researchers, mid-career principal investigators and professors (see full description of the categories in Supporting Information 2). Because the contribution percentages were provided after publication and without discussion among co-authors, these values may not be as accurate as if they had been agreed upon by all co-authors prior to publication. Hence the aim of this exercise was not to produce a highly accurate dataset, and therefore, the following analysis should be regarded as illustrative.

ACI varied from 0.0101, which means the author claims to have produced 101 times less work than his/her co-authors have on average) to 168, which the author claims to have produced 168 times more work than his co-authors have on average (Fig. 4A). Most researchers produced papers with a range of ACI values. Individuals with a majority of high ACI, are likely to be drivers of publications, while those with a majority of medium ACI can be regarded as highly collaborative and those with a majority of low ACI may be service providers. The latter may provide assistance with a limited but potentially essential aspect of the research such as sampling, statistical treatment of the data, supervision or mentoring.

**Figure 4.**
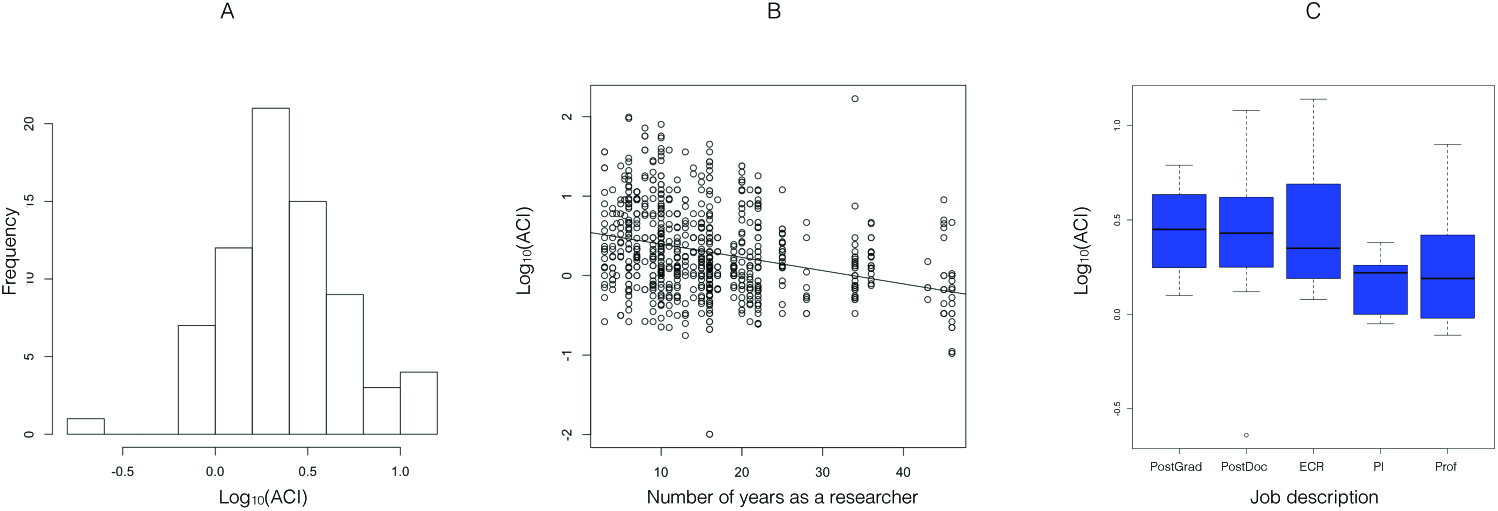
Descriptive statistics of author contribution index (ACI, see Equation 1) for 97 authors between 2014 and 2016 based on an online survey.

ACI will most likely vary across the course of an academic career. In our dataset, ACI varies in relation to the number of years as a researcher (F_1,835_ = 87.57, p<0.0001), however, the correlation remains weak (r=0.306), due to wide variability in ACI (Fig. 4C). With regards to job descriptions, average ACI was higher for non-permanent supervised staff (postgraduate students and postdoctoral fellows) compared to permanent and independent researchers (early career researchers, mid-career principal investigators and Professors) (ANOVA t = 2.327, p = 0.0229) a similar difference was observed when comparing all early career researchers (postgraduate students, postdoc and early career researchers) to established researchers (mid-career principal investigators and Professors) (ANOVA t=2.546, p=0.0132) (Fig. 4). As researchers become independent and establish their own research team, they probably start supervising their own students and postdocs and their ACI is likely to decrease accordingly.

It is possible to link ACI with article impact through the number of citations, altmetrics or any other article-based impact metrics, for example, by dividing ACI by the number of citations for a particular papers. By averaging ACIs, one could also summarise the output of a given scientist as a single number as suggested with other indices (e.g. [4]). However, we do not recommend such practice as it would largely mask the scientist’s output profile, thereby deceiving the purpose of ACI, and it would not be a meaningful way to compare scientist outputs as scientists with very different profiles may reach a very similar average ACI. In our dataset, there are numerous cases in which individuals at very different stages of their career reached a similar average ACI (Fig. 4).

## Conclusion

There are many examples of author contribution indices that have been proposed but none has really been adopted by scientific journals. Many of the proposed solutions are either too complicated, not accurate enough or not comparable across articles, authors and disciplines. The author contribution index presented here addresses these three major issues and if adopted by scientific journals, it could significantly clarify the contribution of co-authors. This index is currently implemented in the recently launched journal *Rethinking Ecology* [3]. We hope that ACI will be adopted by many other journals to increase transparency in co-authored work and attribute accurate credit to authors. Although the current paper uses ecology as the focus, the proposed index is readily applicable to other scientific fields. With regards to past literature and papers published in journals that will not implement ACI, we propose that existing reference list repositories such as Publons (publons.com), ResearchGate (researchgate.net) or ORCID (orcid.org) could provide an option for authors to record the percentage contributions of their publications. Because these values may not be vetted by all co-authors (as opposed to percentage contributions provided at the time of submission), several levels of verification should be displayed for each paper. Values could be considered as 1) unverified if only one co-author provides them, 2) partially verified if at least a second co-author confirms the numbers and 3) fully verified if all co-authors of a paper confirm the numbers. The proposed ACI index has the potential to contribute to more transparency in the science literature it will provide job seekers, recruiters and evaluating bodies with a tool to gather information that is essential to them and cannot be easily and accurately obtained otherwise.

## Material and Methods

The URL address and information about the online survey were distributed through electronic mailing lists, ecological society newsletters and social media. Responses were collected between September 4th 2016 and January 8th 2017. During this timeframe, 97 ecology scientists from 19 different countries completed the survey. Data with contradictory or obviously inaccurate information (for example multi-authored publications where the respondent claims 100% of the work) were removed. The final dataset comprised data for 836 publications from 97 ecology researchers. Author contribution indices (ACIs) of the respondents were calculated for each publications using Equation (1). ACIs were log-transformed (*log*_10_(*ACI)*) to meet the assumptions of normality and all statistical analyses were conducted in R [14]. Respondents were asked to provide information about the number of years they have been research active. This was defined as the time from first year of PhD study or first published peer-reviewed paper, whichever came first. Linear regression and F-statistics were used to analyse ACI in relation to the number of years as an active researcher. Respondents were categorised in different job positions as follow: Postgrad: a postgraduate student; PostDoc: a postdoctoral fellow or other non-permanent staff; ECR: a tenure or permanent early-career researcher; PI: a mid-career principal investigator; Prof: an Associate Professor of Full Professor; Other. ACI was analysed in relation to job description using ANOVA with a priori contrasts between non-permanent supervised staff (postgraduate and postdoc) and permanent and independent researchers (ECR, PI and professors), as well as between all early career researchers (postgraduate, postdoc and ECR) and established researchers (mid-career PI and professors).

## Author contribution

Designed the concept and drafted the manuscript: SB. Prepared the figures: SB, TI. Arranged ethical approval and gathered data from scientists in their respective country: SB,TI, JMO, JS. Reviewed existing journals’ policy: MCL. Contributed to the writing of the final version of the manuscript: SB, TI, JMO, JS, MCL.

## Acknowledgments

We thank the 97 ecologists who responded to the online survey on author contribution.

## Supporting Information

- Supplementary Material 1: Journals policy on authors’ contribution section for the top 150 Ecology journals (according to ISI Web of Knowledge)
- Supplementary Material 2: Co-authorship survey used to collect data from ecology scientists

